# Eukaryotic translation initiation factor 2A protects pancreatic beta cells during endoplasmic reticulum stress while rescuing translation inhibition

**DOI:** 10.1101/2021.02.17.431676

**Authors:** Evgeniy Panzhinskiy, Søs Skovsø, Haoning Howard Cen, Kwan Yi Chu, Kate MacDonald, Galina Soukhatcheva, Derek A. Dionne, Luisa K. Hallmaier-Wacker, Jennifer S. Wildi, Stephanie Marcil, Nilou Noursadeghi, Farnaz Taghizadeh, C. Bruce Verchere, Eric Jan, James D. Johnson

## Abstract

The endoplasmic reticulum (ER) stress-induced unfolded protein response (UPR) helps decide β cell survival in diabetes. The alternative eukaryotic initiation factor 2A (EIF2A) has been proposed to mediate EIF2S1-independent translation during cellular stress and viral infection, but its role in β cells is unknown. EIF2A abundance is high in human and mouse islets relative to other tissues, and both thapsigargin and palmitate significantly increased *EIF2A* mRNA and EIF2A protein levels in MIN6 cells, mouse islets and human islets. Knockdowns of EIF2A, the related factor EIF2D, or both EIF2A and EIF2D, were not sufficient to cause apoptosis. On the other hand, transient or stable EIF2A over-expression protected MIN6 cells, primary mouse islets, and human islets from ER stress-induced, caspase-3-dependent apoptosis. Mechanistically, EIF2A overexpression decreased ERN1 (also known as IRE1α) expression in thapsigargin-treated MIN6 cells or human islets. *In vivo*, β cell specific EIF2A viral overexpression reduced ER stress, improved insulin secretion, and abrogated hyperglycemia in *Ins2*^Akita/WT^ mice. EIF2A overexpression significantly increased expression of genes involved in protein translation and reduced expression of pro-apoptotic genes (e.g. ALDH1A3). Remarkably, the decrease in global protein synthesis during UPR was prevented by EIF2A, despite ER stress-induced EIF2S1 phosphorylation. The protective effects of EIF2A were additive to those of ISRIB, a drug that counteracts the effects of EIF2S1 phosphorylation. Cells overexpressing EIF2A showed higher expression of translation factor EIF2B5, which may contribute to the lack of translational inhibition in these cells. We conclude that EIF2A is a novel target for β cell protection and the circumvention of EIF2S1-mediated translational repression.

## INTRODUCTION

Impaired β cell viability is critical in most, if not all, forms of diabetes (Johnson and Luciani, 2010). Evidence points to a particularly important role for dysregulated protein homeostasis and endoplasmic reticulum (ER) stress as common mechanisms involved in β cell death in type 2 diabetes, type 1 diabetes and islet transplantation (Cnop et al., 2017a; Ghosh et al., 2019). ER stress is managed by signaling pathways collectively called the unfolded protein response (UPR)(Scheuner and Kaufman, 2008). Unfolded proteins are sensed by eukaryotic translation initiation factor 2 α kinase 3 (EIF2AK3, also known as PERK), endoplasmic reticulum to nucleus signaling 1 (ERN1, also known as IRE1α), and activating transcription factor 6 (ATF6). Each of these sensors signals to the nucleus to induce the transcription of stress response genes, including protective ER chaperones and ER-associated degradation proteins to clear misfolded proteins from the ER (Scheuner and Kaufman, 2008). When ER stress becomes unbearable, pro-apoptotic genes are induced, including the master transcription factor DNA damage inducible transcript 3 (DDIT3, also known as CHOP)(Marciniak et al., 2004). Thus, ER stress can stimulate the cell to protect itself, or it can instruct the cell to commit suicide. How β cells make this decision remains incompletely understood and mechanistic detail is required for therapeutic interventions to prevent diabetes.

Attenuation of global protein translational in response to ER stress prevents the further translation and accumulation of unfolded proteins (Cnop et al., 2017b). Paradoxically, although ER stress leads to translational inhibition by phosphorylated EIF2S1 (also known as EIF2α), a subset of cellular and viral mRNAs is translated despite EIF2S1 phosphorylation. mRNAs important for stress amelioration are preferentially translated through upstream open reading frame-mediated mechanisms during activation of the ER stress (Starck et al., 2016), but the mechanisms underlying selective translation have not been clearly established (Hinnebusch and Lorsch, 2012; Lee et al., 2009; Young and Wek, 2016). In the cases of ATF4, ATF5, and GADD34, this involves upstream open reading frame-initiated translation (Hinnebusch and Lorsch, 2012; Lee et al., 2009). However, translation factors involved in this and other types of non-canonical initiation of protein synthesis are still poorly understood.

One candidate factor that mediates non-canonical translation initiation, called eukaryotic initiation factor 2A (EIF2A), can stimulate binding of initiator methionine tRNA (Met-tRNA^Met^_i_) to 40S ribosomal subunits *in vitro* (Adams et al., 1975; Merrick and Anderson, 1975). EIF2A should not be confused with the α subunit of the eukaryotic initiation factor 2 (official gene name EIF2S1), which is responsible for the binding of Met-tRNA ^Met^_i_ to 40S ribosomal subunits in GTP dependent manner (Benne et al., 1979; Staehelin et al., 1979). During ER stress, EIF2AK3 reduces the level of active EIF2 by phosphorylating EIF2S1, consequently, reducing global translation (Back et al., 2009; Donnelly et al., 2013). In contrast to EIF2, EIF2A directs binding of the Met-tRNA^Met^_i_ to 40S ribosomal subunits in a codon-dependent manner (Zoll et al., 2002). In yeast, EIF2A has been implicated in internal ribosome entry site-mediated translation (Reineke et al., 2011; Zoll et al., 2002). In mammalian cells, EIF2A helps viral mRNAs avoid inhibition of translation, when double-stranded RNA-activated EIF2AK2 phosphorylates EIF2S1 (Kim et al., 2011; Ventoso et al., 2006). Translation of a few endogenous mRNAs are EIF2A-dependent (Liang et al., 2014; Starck et al., 2012; Starck et al., 2016), but it does not appear that this factor plays a non-redundant role in translation since EIF2A-total knockout mice are viable (Golovko et al., 2016). It is plausible that other related factors, such as EIF2D (also known as ligatin)(Dmitriev et al., 2010) or EIF5B (Kim et al., 2018; Wang et al., 2019) may compensate for the EIF2A loss-of-function. Targeting alternative translation initiation represents a promising therapeutic strategy. For example, the experimental drug ISRIB (integrated stress response inhibitor) allows bypass of translation initiation attenuation induced by EIF2S1 phosphorylation (Rabouw et al., 2019) and is in clinical trials for multiple conditions where ER stress is a contributing factor.

In the present study, we investigated involvement of alternative translation initiation factor EIF2A in β cell UPR *in vitro* and *in vivo*. We showed that EIF2A was highly expressed in β cells and was upregulated during ER stress. Overexpression of EIF2A *in vitro* attenuated ER stress induced apoptosis in β cells. We also demonstrated that overexpression of eIF2A in pancreatic β cells prevented progression of diabetes and potentiates glucose stimulated insulin secretion in *Ins2*^Akita/WT^ mice. EIF2A drove a protective gene expression profile and circumvented the phospho-EIF2S1 block on protein translation in manner distinct from ISRIB.

## RESULTS

### EIF2A is highly expressed in pancreatic β cells and induced during ER stress

We started by examining the protein abundance and mRNA expression of EIF2A and EIF2D across human tissues, and compared them with the canonical initiation factor EIF2S1. Publicly available mass spec (www.proteomicsdb.org) and RNAseq data (Human Cell Atlas) show that EIF2A is differentially expressed between tissues, with the highest protein levels present in pancreatic islets. The elevated islet-specific protein abundance of EIF2A was driven, at least in part, by relatively high mRNA expression (Fig. 1A). EIF2D was not highly expressed under basal conditions and was not enriched in islets compared to other tissues (Fig. 1A). EIF2S1 is highly and evenly abundant at the protein level across analyzed tissues despite some differences in mRNA expression (Fig. 1A). To extend these findings to mice, we quantified protein levels of EIF2A, EIF2D, and EIF2S1 across different tissues in C57BL/6/J mice using Western Blot analysis. Consistent with the human data, pancreatic islets had highest EIF2A protein abundance compared to the other analyzed tissues. EIF2D protein levels in pancreatic islets were similar to spleen, muscle and liver, but significantly higher than in brain and heart (Fig. 1B). We then analyzed abundance of translation initiation factors in human islets isolated from donors with and without type 2 diabetes. Islets from diabetic donors had higher levels of phosphorylated EIF2S1 indicating elevated ER stress levels, however abundance of EIF2S1 and EIF2A was not significantly affected by diabetes status (Fig. 1C). Interestingly, we found a positive correlation between EIF2A and EIF2S1 protein levels in human islets from donors with type 2 diabetes (Fig. 1C). Islets of Langerhans contain multiple cell types, so in order to identify which cells express these alternative translation initiation factors we used immunohistochemistry staining. EIF2A and EIF2D were primarily co-localized with insulin positive β cells in mouse pancreas sections (Fig. 1D, E). EIF2A was localized to the endoplasmic reticulum of the MIN6 mouse insulinoma β cell line (Fig. 1F). Together, these observations indicated that EIF2A is abundant in β cells and correlates with ER stress in T2D human islets.

**Figure 1.**
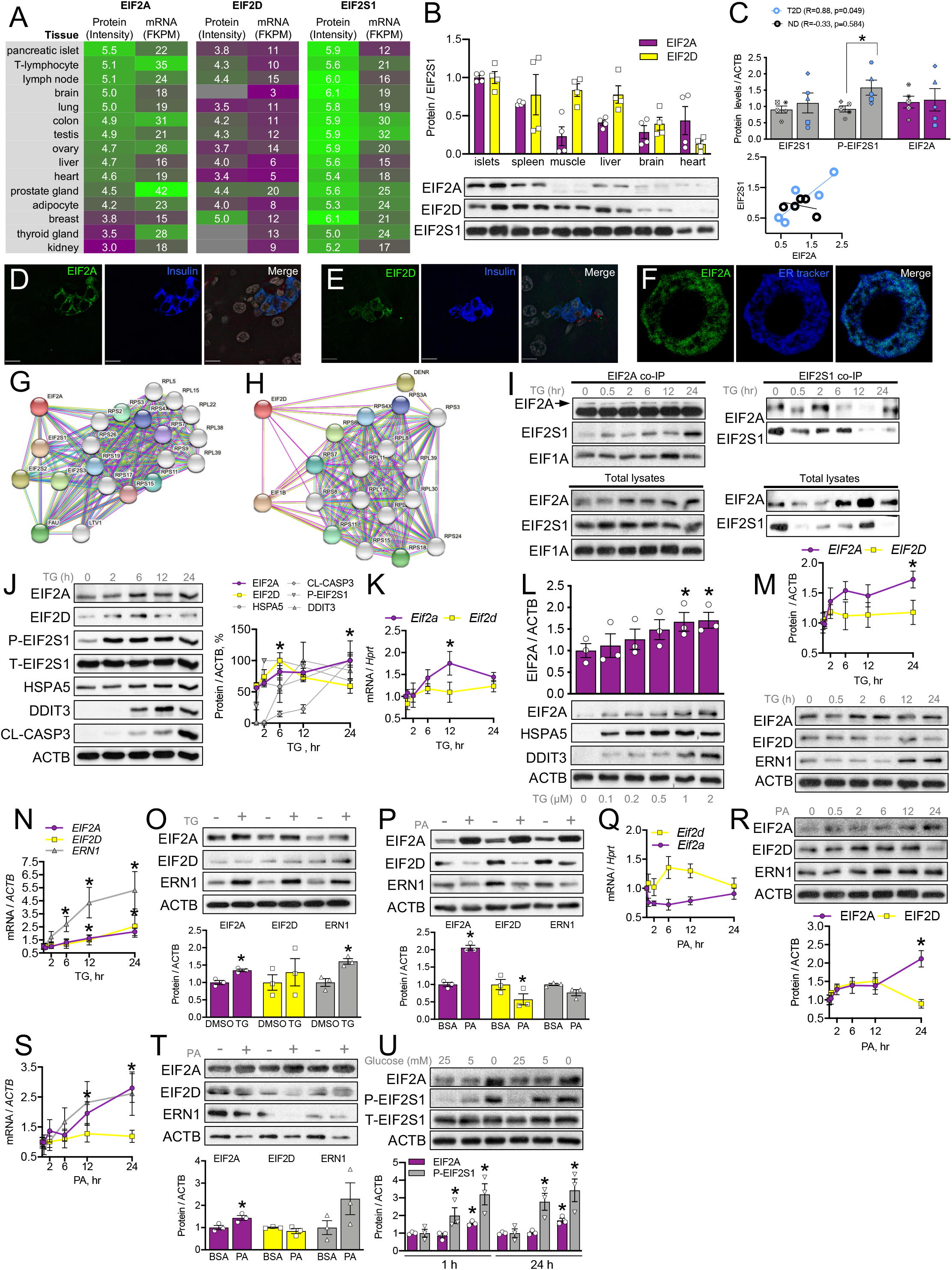
Islet enriched EIF2A is induced by ER stress in β cells. **(A)** Protein abundance and mRNA expression of EIF2A, EIF2D and EIF2S1 across different human tissues. Data from human tissue proteomics (www.proteomicsdb.org) and RNAseq (HCA). **(B)** EIF2A, EIF2D and EIF2S1 protein abundance in mouse tissues (n=4). **(C)** Correlation between and abundance of EIF2A and EIF2S1 protein levels in human islets from non-diabetic and type 2 diabetic donors (n=5). **(D**,**E)** EIF2A and EIF2D proteins are enriched in mouse islets relative to other pancreatic cell types *in vivo*. **(F)** EIF2A is an ER-resident protein in MIN6 cells. **(G**,**H)** Predictions of physical network interactions for EIF2A **(G)** and EIF2D **(H)** with STRING v11, confidence level = 0.7. **(I)** Upper panels: representative immunoblots of lysates using antibodies against EIF2A, EIF2S1, EIF1A and EIF5A after immunoprecipitation using anti-EIF2A (left panel) or anti-EIF2S1antibodies (right panel). Bottom panel: non-precipitated total lysates control. **(J)** Representative immunoblots and densitometry analysis of protein levels of EIF2A and ER stress markers relative to ACTB in MIN6 cells treated with 1 μM thapsigargin (TG) for 0, 2, 6, 12, and 24 hours (n=3-4). **(K)** Quantification of *Eif2a* and *Eif2d* mRNA levels relative to *Hprt* in MIN6 cells treated with 1 μM thapsigargin (TG) for 0, 0.5, 2, 6, 12, and 24 hours (n=3-4). **(L)** Representative immunoblots and densitometry quantification of EIF2A expression in dose response to 0, 0.1, 0.2, .05, 1 and 2 μM thapsigargin (TG) treatment for 24 hours in MIN6 cells (n=3). **(M**,**N)** Representative immunoblots and densitometry **(M)** and mRNA levels **(N)** of *EIF2A, EIF2D* and *ERN1* relative to ACTB in dispersed human islets treated with 1 μM thapsigargin (TG) for 0, 0.5, 2, 6, 12 or 24 hours. **(O**,**P)** Representative immunoblots and densitometry quantification of EIF2A, EIF2D and ERN1 expression in **(O)** isolated mouse islets treated with 1 μM thapsigargin (TG) for 24 hours or **(P)** MIN6 cells treated with 0.5 mM palmitate (PA) for 24 hours. **(Q)** Quantification of *Eif2a* and *Eif2d* mRNA levels relative to *Hprt* in MIN6 cells treated with 0.5 mM palmitate (PA) for 0, 0.5, 2, 6, 12, and 24 hours (n=3-4). **(R**,**S)** Representative immunoblots and densitometry **(R)** and mRNA levels **(S)** of *EIF2A, EIF2D* and *ERN1* relative to *ACTB* in dispersed human islets treated with 0.5 mM palmitate (PA) for 0, 0.5, 2, 6, 12 or 24 hours (n=3). **(T)** Representative immunoblots and densitometry quantification of EIF2A, EIF2D and ERN1 expression in isolated mouse islets treated with with 0.5 mM palmitate (PA) for 24 hours. **(U)** Representative immunoblots and densitometry quantification of EIF2A protein and EIF2S1 phosphorylation levels in response to 0 mM, 5 mM or 25 mM glucose in MIN6 cells. *p<0.05 compared to vehicle treated cells.

We used STRING software to predict physical interaction human protein networks involving EIF2A and EIF2D. With high confidence, EIF2A was predicted to bind EIF2S1, EIF2S2 and EIF2S3, as well as multiple ribosomal proteins (Fig. 1G). EIF2D was predicated to bind EIF1B, DENR (another alternative translation factor) and multiple ribosomal proteins (Fig. 1H). Using co-IP and reverse co-IP, we confirmed that EIF2A physically interacts with EIF2S1 in β cells, in an ER-stress independent manner (Fig. 1I). These bioinformatic and biochemical analyses illustrate that EIF2A and EIF2D are integral to the cellular protein synthesis machinery.

We analyzed expression of these alternative translation initiation factors during UPR in β cells, when canonical translation is inhibited. We used the established ER stress inducer thapsigargin, which decreases ER calcium levels by inhibiting sarcoplasmic/endoplasmic reticulum Ca^2+^-ATPases (Gwiazda et al., 2009). As expected, EIF2S1 was phosphorylated within 2 hours of thapsigargin treatment. EIF2S1 phosphorylation was sustained up to 24 hours of thapsigargin treatment. Thapsigargin significantly increased protein and mRNA expression of EIF2A in MIN6 cells, isolated mouse or human islets (Fig.1J, K, M, N, O). We observed the highest levels of EIF2A protein 24 hours after induction of UPR, which coincided with the highest induction of the ER stress markers heat shock protein A5 (HSPA5, also known as GRP78 or BiP), DDIT3 and ERN1. EIF2A protein up-regulation in response to thapsigargin treatment was dose dependent (Fig. 1L). EIF2D showed only transient increase on a protein level in MIN6 cells following ER stress induction with thapsigargin, but not in isolated mouse or human islets (Fig. 1J-O).

We also employed another ER stress inducer, palmitate, which mimics lipotoxicity-induced ER stress observed in type 2 diabetes *in vitro* (Gwiazda et al., 2009). Palmitate treatment increased EIF2A protein (Fig. 1P) but not mRNA levels (Fig. 1Q) in MIN6 cells. On the other hand, EIF2D protein was downregulated after 24 hours of palmitate exposure (Fig. 1P). We validated these findings in isolated human islets, showing that palmitate induced EIF2A protein and mRNA levels, but not EIF2D protein and mRNA levels (Fig. 1R, S). In isolated primary mouse pancreatic islets palmitate exposure for 24 hours induced EIF2A protein, but not EIF2D protein (Fig. 1T). Together, these results demonstrate a strong association between UPR and EIF2A levels.

We also tested whether EIF2A expression was induced under other conditions that induce EIF2S1 phosphorylation in β cells, such as low glucose concentration (Cavener et al., 2010). EIF2A protein expression was significantly increased after 1 hour of glucose deprivation, which coincided with increase in EIF2S1 phosphorylation in MIN6 cells (Fig. 1U). This effect was still observed 24 hours after glucose withdrawal (Fig. 1U).

### EIF2A knockdown leads to EIF2D upregulation during ER stress-induced apoptosis

Induction of EIF2A expression during UPR in β cells prompted us to investigate effect of EIF2A loss-of-function on β cell survival during ER stress. We successfully reduced EIF2A protein abundance in MIN6 cells using shRNA-mediated knockdown and prevented increased EIF2A expression in response to thapsigargin treatment (Fig. 2A). However, MIN6 cells with reduced EIF2A levels had the same level of ER stress-induced apoptosis after thapsigargin treatment (Fig. 2B). Compared with MIN6 cells containing the scramble control, EIF2A knockdown resulted in no difference in induction of phosphorylation of EIF2S1, or expression of HSPA5 and DDIT3, indicating a normal UPR response (Fig. 2C). However, EIF2A knockdown led to a significant increase in abundance of another alternative translation initiation factor EIF2D, suggesting compensatory mechanisms and functional redundancy between EIF2A and EIF2D (Fig. 2C).

**Figure 2.**
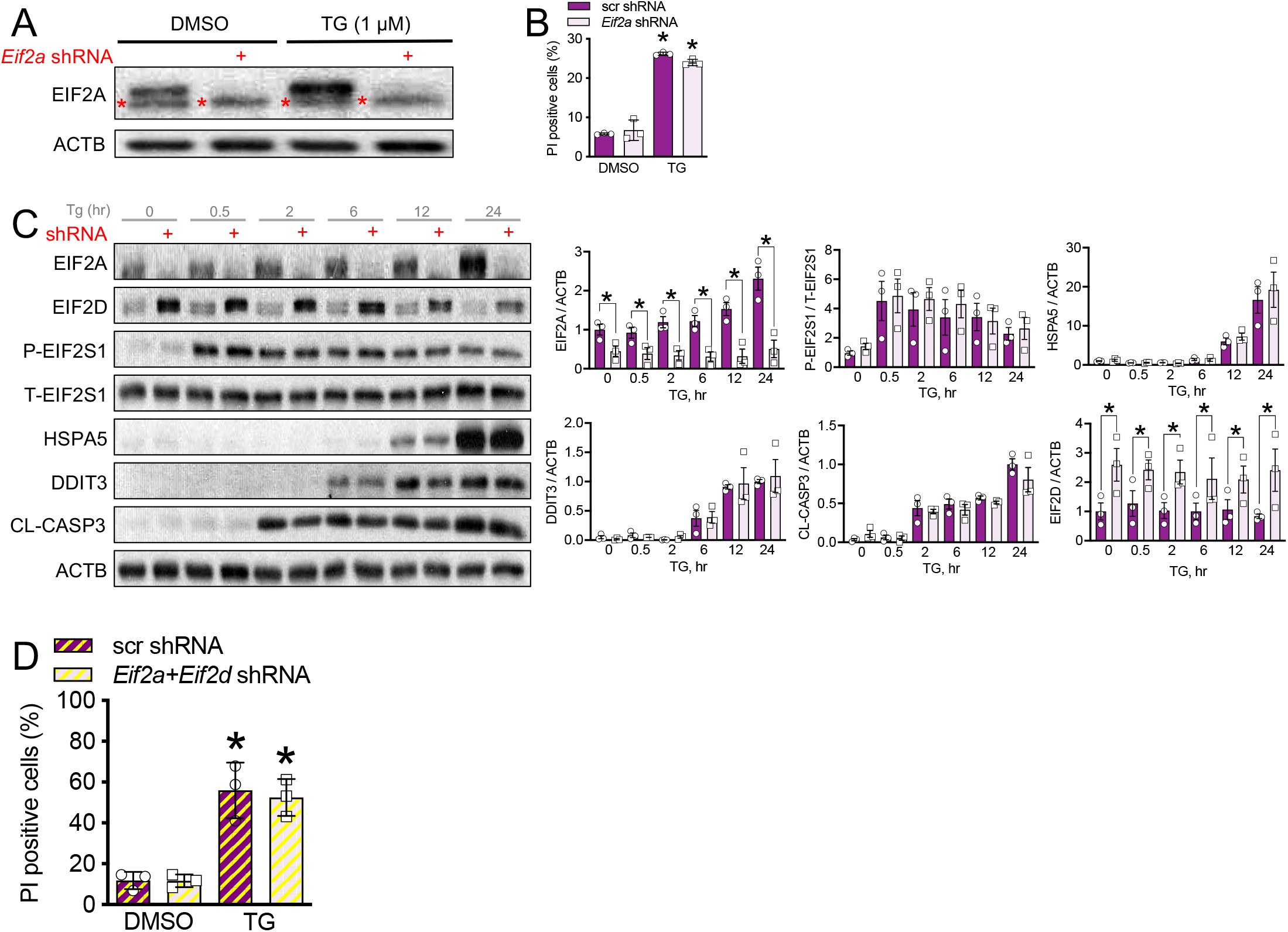
Effects of EIF2A knockdown in MIN6 cells. **(A)** Representative immunoblots confirming knockdown of EIF2A in MIN6 cells in the presence of 1 μM thapsigargin (TG). **(B)** Quantification of propidium iodide (PI) cell death assay in MIN6 cells transfected with scramble control (scr shRA) or shRNA targeting *Eif2a* and treated with 1 μM thapsigargin (TG) or DMSO for 24 hours (n=3), *p<0.05 compared to 0.1% DMSO treated cells. **(C)** Representative immunoblots and densitometry quantification of protein abundance of EIF2A, phosphorylated EIF2S1, HSPA5, DDIT3, cleaved caspase 3 and EIF2D relative to ACTB in MIN6 cells transfected with scramble shRNA or shRNA targeting *Eif2a* and treated with 1 μM thapsigargin for 0, 0.5, 2, 6, 12 or 24 hours (n=3). **(D)** Quantification of propidium iodide (PI) cell death assay in MIN6 cells transfected with scramble shRNA (scr shRNA) or shRNAs targeting *Eif2a* and *Eif2d* (*Eif2a*+*Eif2d* shRNA) and treated with 1 μM thapsigargin or 0.1% DMSO for 48 hours (n=3), *p<0.05 compared to 0.1% DMSO treated cells

We performed EIF2A/EIF2D double knockdown to determine whether ER stress or ER stress-induced apoptosis might be exacerbated. Despite the successful reduction of both EIF2A and EIF2D protein levels, we did not observe any significant effects on ER stress induced apoptosis in MIN6 cells. Phosphorylation of EIF2S1 and expression of pro-apoptotic DDIT3 during UPR were also unaffected by EIF2A/EIF2D double knockdown. Since EIF2A and EIF2D are not the only proteins that can promote efficient recruitment of Met-tRNA^Met^_i_ under conditions of inhibition of EIF2 activity (Schleich et al., 2014; Skabkin et al., 2010; Wong et al., 2018), it is possible that there are multiple levels of redundancy.

### EIF2A overexpression protects pancreatic β cells from ER stress-induced apoptosis

Given that endogenous expression of EIF2A is increased during ER stress, we tested whether EIF2A overexpression might be sufficient to protect β cells under stress conditions. Strikingly, lentiviral infection to achieve a 3-fold increase in MIN6 cells (Fig. 3A) resulted in significant reductions in thapsigargin-(Fig. 3B) and palmitate-induced apoptosis (Fig. 3K), compared to cells overexpressing GFP only. The reduced cell death by stable EIF2A overexpression in MIN6 cells was associated with sustained insulin secretion during UPR (Fig. 3C). We analyzed mRNA levels of common UPR markers (Fig. 3D-G) and observed significant decrease in *Ern1* expression (Fig. 3G) after 24 hr of thapsigargin treatment in MIN6 cells overexpressing EIF2A, suggesting selective inhibition of the ERN1 branch of UPR upon increased activity of EIF2A. We also observed a transient increase in *Hspa5* mRNA levels in EIF2A-overexpressing cells after 6 hours of thapsigargin treatment (Fig. 3D). We showed significant decrease in protein expression of UPR pro-apoptotic markers DDIT3 and inhibition of caspase 3 activation in EIF2A-overexpressing cells. However, there was no effect on the steady-state levels of *Ddit3* mRNA during UPR (Fig. 3E), suggesting a specific role of EIF2A in the inhibition of ER stress-induced translation of *Ddit3* mRNA. Interestingly, DDIT3 translation may be regulated by an upstream open reading frame (Chen et al., 2010). There was no change in protein expression of molecular chaperone HSPA5 or in translation-inhibiting phosphorylation of EIF2S1 (Fig. 3H).

**Figure 3.**
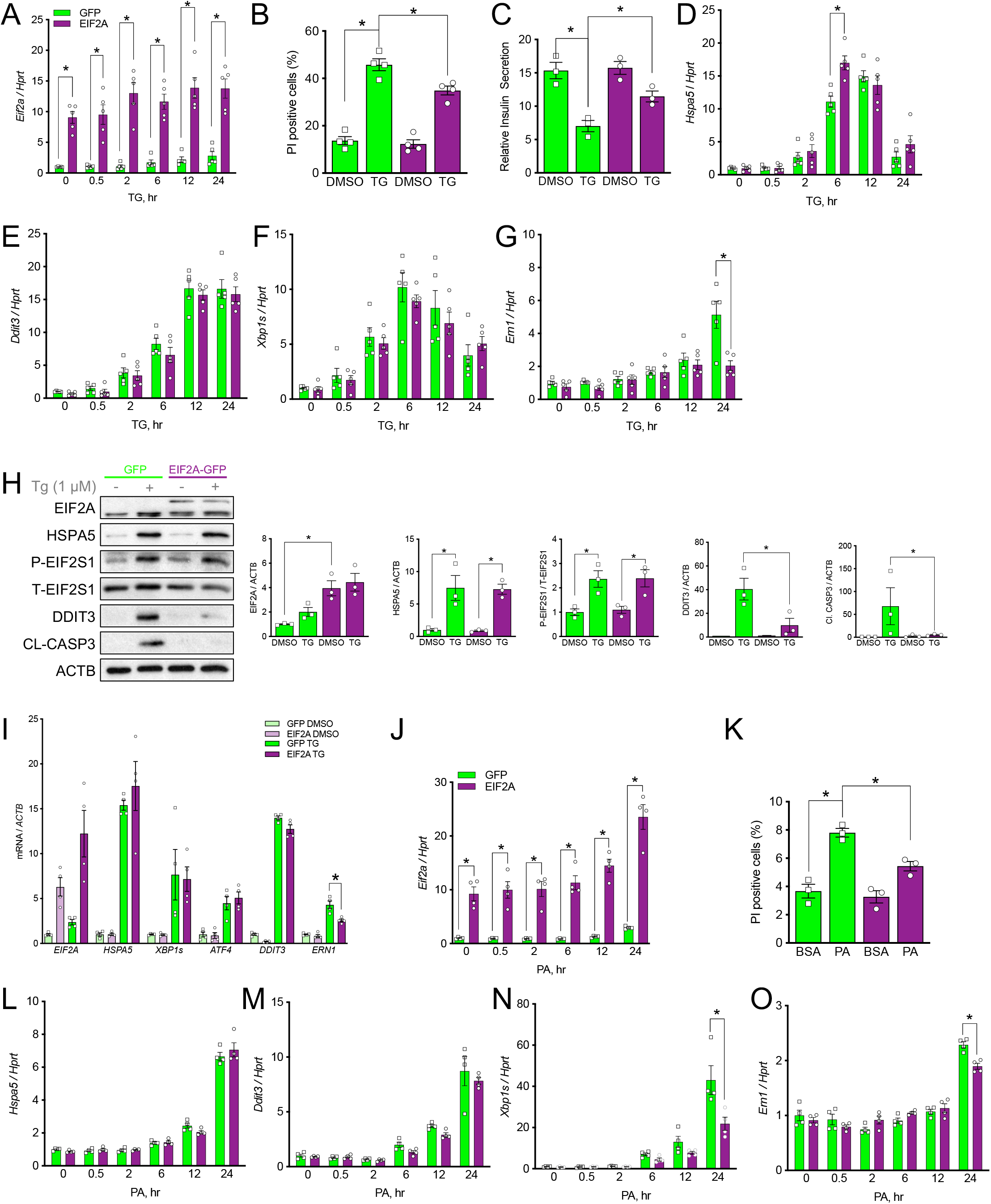
EIF2A over-expression protects MIN6 cells and human islets from ER stress-induced apoptosis. **(A)** qPCR confirming overexpression of *Eif2a* relative to *Hprt* in MIN6 cells in the presence of 1 μM thapsigargin (TG) for 0, 0.5, 2, 6, 12 or 24 hours. **(B)** Quantification of propidium iodide (PI) cell death assay in MIN6 cells transfected with a plasmid expressing GFP or EIF2A-GFP and treated with 1 μM thapsigargin (TG). (**C)** Static insulin secretion assay in MIN6 cells overexpressing EIF2A-GFP or GFP and treated with thapsigargin (TG) for 24 hours. **(D-G)** mRNA levels of *Hspa5* **(D)**, *Ddit3* **(E)**, spliced *Xbp1* **(F)**, and *Ern1* **(G)** relative to *Hprt* in MIN6 cells transfected with a plasmid expressing GFP or EIF2A-GFP and treated with thapsigargin for 0, 0.5, 2, 6, 12 or 24 hours. **(H)** Representative immunoblots and densitometry quantification expression of EIF2A, HSPA5, phosphorylated EIF2S1, DDIT3, and cleaved caspase 3 relative to ACTB in MIN6 cells transfected with a plasmid expressing GFP or EIF2A-GFP and treated with DMSO or thapsigargin for 24 hours. **(I)** mRNA levels of *EIF2A, HSPA5*, spliced *XBP1* (*XBP1s*), *ATF4, DDIT3* and *ERN1* relative to *HPRT* in dispersed human islets infected with a lentivirus expressing GFP or EIF2A-GFP and treated with thapsigargin for 24 hours. **(J)** qPCR confirming overexpression of *Eif2a* relative to *Hprt* in MIN6 cells in the presence of 0.5 mM palmitate (PA) for 0, 0.5, 2, 6, 12 or 24 hours. **(K)** Quantification of propidium iodide (PI) cell death assay in MIN6 cells transfected with a plasmid expressing GFP or EIF2A-GFP and treated with 0.5 mM palmitate (PA). **(L-O)** mRNA levels of *Hspa5* **(L)**, *Ddit3* **(M)**, spliced *Xbp1* **(N)**, and *Ern1* **(O)** relative to *Hprt* in MIN6 cells transfected with a plasmid expressing GFP or EIF2A-GFP and treated with 0.5 mM palmitate (PA). *p<0.05

We validated these findings in isolated human islets, showing that EIF2A overexpression selectively inhibited ERN1 branch of UPR upon increased expression of EIF2A in thapsigargin treated islets (Fig. 3I). Again, we employed another ER stress inducer palmitate to see if the observed molecular effects are specific to the ER inducer or if it is a general response to UPR. Similar to thapsigargin treated cells we observed significant decrease in *Xbp1* splicing (Fig. 3N) and *Ern1* expression (Fig. 3O) in EIF2A-overexpressing MIN6 cells treated with palmitate for 24 hours compared to GFP-overexpressing cells, with no difference in *Hspa5* (Fig. 3L) and *Ddit3* (Fig. 3M) mRNA levels. Taken together, our *in vitro* findings clearly demonstrate that EIF2A protects MIN6 cells and human islets from ER stress-induced apoptosis.

### Overexpression of EIF2A in pancreatic β cells attenuates diabetes progression in Akita mice

To confirm the protective effect of EIF2A overexpression from ER stress-induced apoptosis *in vivo*, we used C57BL/6-*Ins2*^Akita^/J (Akita) mice, a strain that spontaneously develops hyperglycemia due to reduced β cell mass, but without insulitis or obesity (Yoshioka et al., 1997). Diabetic phenotype of these mice is caused by autosomal dominant *Ins2* mutation that cannot fold properly leading to β cell specific ER stress-induced apoptosis (Kayo and Koizumi, 1998; Ron, 2002). Homozygous *Ins2* Akita mutation leads to perinatal lethality; heterozygous mice are viable and fertile, but develop hyperglycemia around 3–4 weeks of age (Chang and Gurley, 2012). Also increase in blood glucose levels is more rapid in male than in female mice (Oyadomari et al., 2002). Therefore, we used heterozygous female *Ins2*^Akita/WT^ mice for our *in vivo* studies, because maximum expression of transgene delivered with adeno-associated virus (AAV) is observed 3 weeks post transduction (Quirin et al., 2018) (Fig. 4C). For β cell specific overexpression, we designed adeno-associated virus 6 (AAV6), encoding either EIF2A-GFP or control GFP and driven by a rat *Ins2* gene promoter (Fig. 4A) and delivered by intraductal injection followed by 4 weeks of monitoring (Fig. 4B). We chose local intraductal delivery of AAV6 for β cell specific overexpression, because it was demonstrated to be quick, efficient and safe way for widespread stable β cell gene transfer (Quirin et al., 2018) (Fig. 4C, M). As expected, *Ins2*^Akita/WT^ mice given GFP-control AAV6 showed increased fasting blood glucose levels with age, but this increase was significantly attenuated by EIF2A overexpression in β cells (Fig. 4E). *Ins2*^Akita/WT^ mice with EIF2A overexpression had lower fasting blood glucose levels compared to GFP control mice at 2 weeks and 3 weeks after AAV6 ductal injections (Fig. 4E). Specific overexpression of EIF2A in β cells had no effect on body weight gain at any time point when compared to GFP controls (Fig. 4D). Three weeks after viral injection, *Ins2*^Akita/WT^ mice overexpressing β cell specific EIF2A showed significantly improved glucose tolerance when compared with controls transduced with AAV6-GFP alone (Fig. 4F). Overexpression of EIF2A in *Ins2*^Akita/WT^ mice was associated with increased insulin secretion at 15 min post glucose challenge (Fig. 4G). In the random fed state, no significant differences in plasma insulin or proinsulin levels were detected between EIF2A-GFP and control GFP groups (Fig. 4J). We assessed *in vivo* the mechanisms associated with improved insulin secretion and glucose homeostasis in AAV6-EIF2A treated *Ins2*^Akita/WT^ mice. In the relatively small number of mice studied, we were unable to observe a statistically significant increase in β cell area (Fig. 4H) or any changes in mRNA expression of *Ins1* or *Ins2* genes (Fig. 4I). However, qPCR analysis revealed significantly reduced expression of the pro-apoptotic ER stress marker *Ddit3* in islets isolated from EIF2A-overexpressing *Ins2*^Akita/WT^ mice (Fig. 4K), with no changes in ER stress-independent apoptotic markers (Fig. 4L). At the protein level, pancreatic islets isolated from *Ins2*^Akita/WT^ mice overexpressing EIF2A showed decreased phosphorylation of EIF2S1 (Fig. 4M). Additionally, β cells in *Ins2*^Akita/WT^ mice overexpressing EIF2A exhibited increase in intensity of immunofluorescent staining for pro-survival molecular chaperone HSPA5 when compared to controls (Fig. 4N), but no change in intensity for pro-apoptotic ER stress marker DDIT3 (Fig. 4O). Collectively, these *in vivo* data support our *in vitro* experiments that illustrated powerful pro-survival effects of modest EIF2A over-expression.

**Figure 4.**
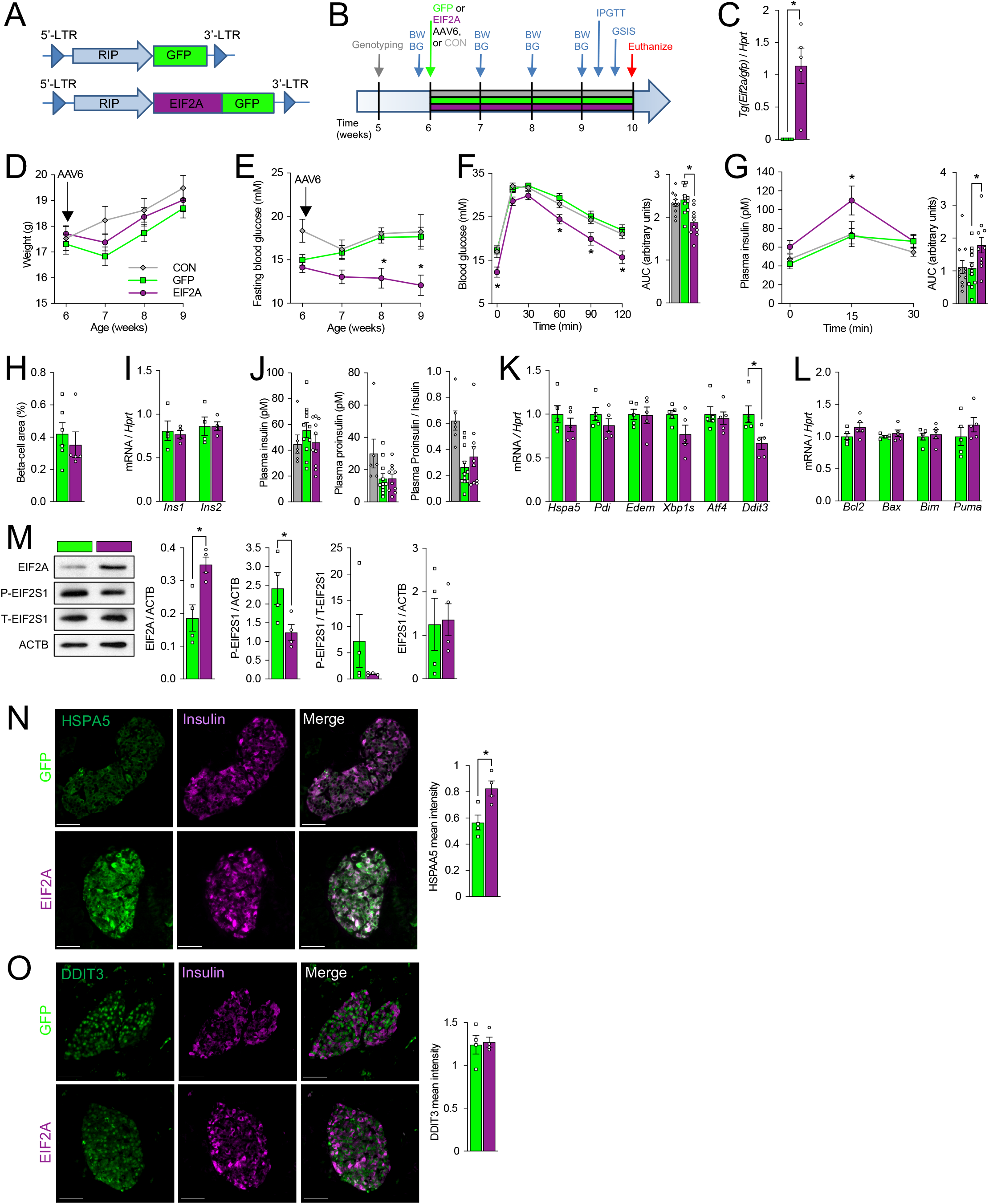
Over-expression of EIF2A by intraductal pancreas AAV-6 injection in 6 wk female C57BL/6J-Ins2Akita/WT mice reversed hyperglycemia, improved glucose tolerance and insulin secretion in Akita mice. **(A)** AAV constructs and **(B)** timeline for *in vivo* experiments. **(C)** Expression of EIF2A:GFP transgene in islets isolated from experimental vs control mice 4 weeks post AAV-6 injection. **(D)** Body weights and **(E)** fasting blood glucose levels in mice following the surgery (n=10-11). **(F)** Blood glucose measurements following IPGTT (1g/kg) and Area Under the Curve (AUC) calculations 3 weeks post intraductal AAV-6 injection (n=10-11). **(G)** *In vivo* GSIS and Area Under the Curve (AUC) calculations 3.5 weeks post intraductal AAV-6 injection (n=10-11). **(H)** Quantification of intensity of immunofluorescence in pancreatic sections stained against insulin. **(I)** *Ins1* and *Ins2* expression in islets isolated from mice 4 weeks post AAV-6 injection. **(J)** Plasma levels of insulin, proinsulin and proinsulin to insulin ratio 3.5 weeks’ post intraductal AAV-6 injection. **(K**,**L)** qPCR quantification of mRNA levels of ER stress markers **(K)** or apoptosis markers **(L)** in islets isolated from mice 4 weeks post AAV-6 injection. **(M)** Representative immunoblots and densitometry quantification of protein levels of HSPA5, EIF2A and (P)-EIF2S1 relative to ACTB or total(T)-EIF2S1 in mouse islets isolated 4 weeks post AAV-6 injection. **(N, O)** Immunofluorescence staining and corresponding quantification of pancreatic sections stained against: insulin and HSPA5 **(N)**, or insulin and DDIT3 **(O)**. *p<0.05

### EIF2A overexpression reprograms the β cell transcriptome

We performed RNAseq to investigate in an unbiased manner the global transcriptomic changes induced by EIF2A overexpression in β cells. MIN6 cells overexpressing EIF2A or GFP were treated for 1 or 3 hours with thapsigargin or DMSO control, and their transcriptomes were analyzed (Fig. 5A). Aldehyde dehydrogenase 1a3 (*Aldh1a3*), which defines a subset of failing pancreatic β cells, was significantly decreased in MIN6 cells overexpressing EIF2A (Kim-Muller et al., 2016). Among genes significantly upregulated in MIN6 cells over-expressing EIF2A were alternative translation initiation factor *Denr* (Schleich et al., 2014), transcriptional activator of glucose-stimulated insulin secretion anoctamin 1 (*Ano1*), (Crutzen et al., 2016; Edlund et al., 2014; Jian and Felsenfeld, 2018), and *Ins1* gene itself. Protein-protein interaction network mapping illustrated a coordinated increase in interaction of ribosomal, proteasomal, and mitochondrial genes, and well-connected hubs including *Crebbp, Hsp90aa1*, and *Hras* (Fig. 5B). There was a decrease in the expression of well-connect hubs *Ep300* and *Foxo3* (Fig. 5B). Pathway analysis identified significantly over-represented gene categories including Ribosome, Huntington’s disease, Parkinson’s disease, and Oxidative Phosphorylation; these effects were especially pronounced after 1 hour of thapsigargin-induced ER stress (Fig. 5C). Using the list of differentially expressed genes, we also predicted the core transcription factor network mediating transcriptomic differences between control and EIF2A-overexpressing cells (Fig. 5D), identifying *Ubtf, Hcfc1, Zmiz1, Mxi1*, and *Myb* as the 5 most connected transcription factors upstream of our differentially expressed genes. Ubtf controls ribosome expression (Edvardson et al., 2017), Hcfc1 is a E2f1 co-activator controlling the expression of the β cell master transcriptional regulator *Pdx1* (Iwata et al., 2013), and Zmiz1 is a positive regulator of β cell function (van de Bunt et al., 2015). Together, these data reveal the molecular mechanisms by which EIF2A protects β cells and suggest a role in global translational regulation.

**Figure 5.**
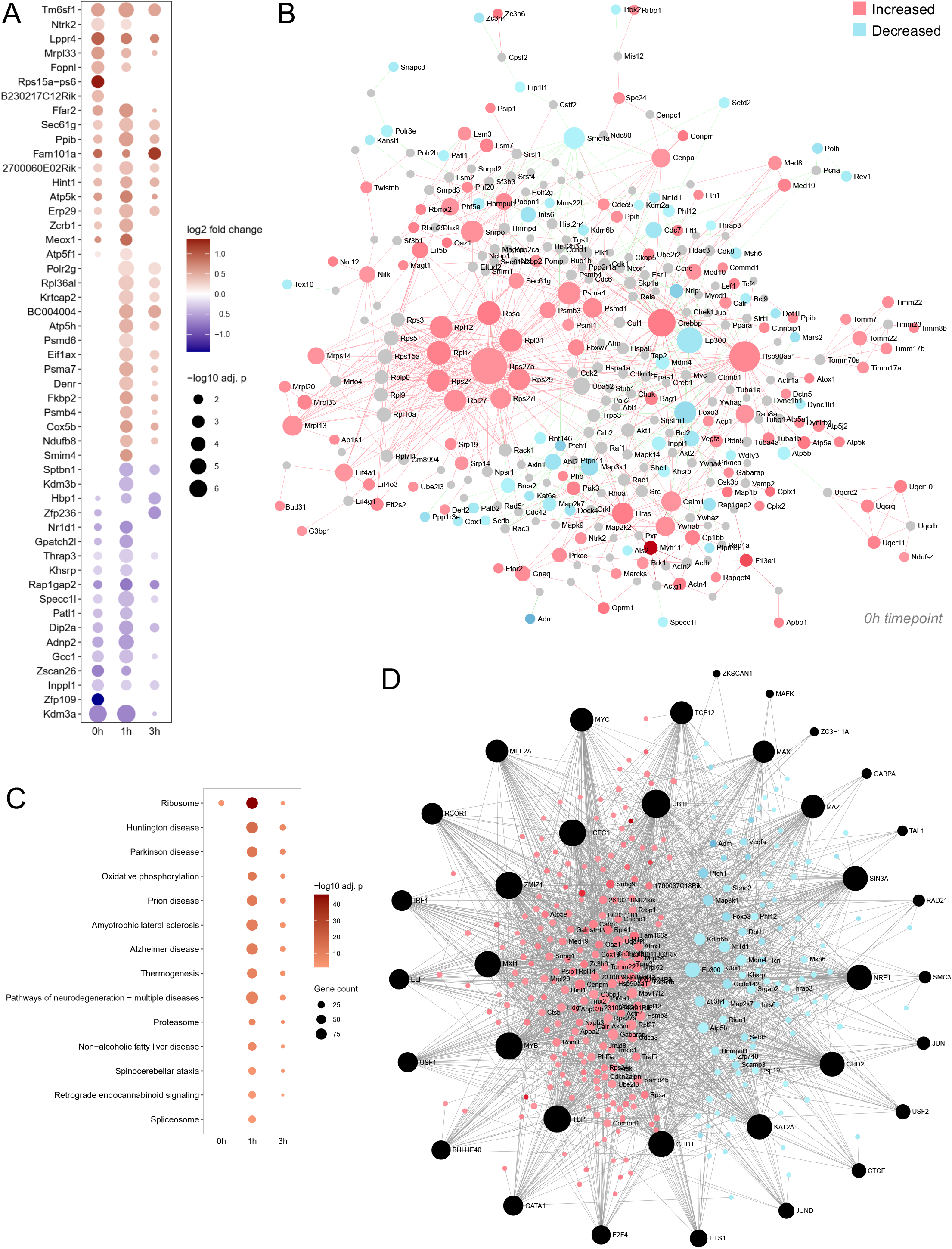
Transcriptional consequences of EIF2A:GFP over-expression in the context of ER stress. **(A)** Fold change of the top 50 differentially expressed genes from MIN6 cells overexpressing GFP (control) or EIF2A:GFP and treated with 1 μM thapsigargin for 0 (DMSO control), 1 or 3 hours. **(B)** Protein-protein interaction network of all the differentially expressed genes between MIN6 cells overexpressing GFP (control) and EIF2A:GFP. **(C)** KEGG pathways enriched from differentially expressed genes between MIN6 cells overexpressing GFP (control) and EIF2A:GFP from 0 (0.1% DMSO control), 1 or 3-hour 1 μM thapsigargin treatment. **(D)** Upstream transcription factors predicted from differentially expressed genes between MIN6 cells overexpressing GFP (control) and EIF2A:GFP.

### EIF2A overexpression reverses the ER stress-induced block on protein synthesis

A hallmark of the UPR is the transient attenuation of global protein synthesis (Cnop et al., 2017a; Ghosh et al., 2019). In our hands, MIN6 cells treated with thapsigargin exhibit a rapid (30 minutes) and sustained (at least 20 hours) decreased protein synthesis rate by 80.2 ± 10% (Fig. 6A). In contrast to other cells types such as hepatocytes, global protein synthesis rates did not significantly recover under these conditions and EIF2S1 remained phosphorylated in β cells (Fig. 1J). We originally wondered whether EIF2A may facilitate the translation of specific mRNAs and we were therefore surprised to find that EIF2A overexpression was sufficient to completely prevent thapsigargin-induced translation inhibition (Fig. 6A). We confirmed that EIF2A overexpression also prevented the loss of high molecular weight polysomes in thapsigargin-treated β-cells (Fig. 6B). These data demonstrate that elevated EIF2A circumvents the block on protein translation induced by phosphorylated EIF2S1.

**Figure 6.**
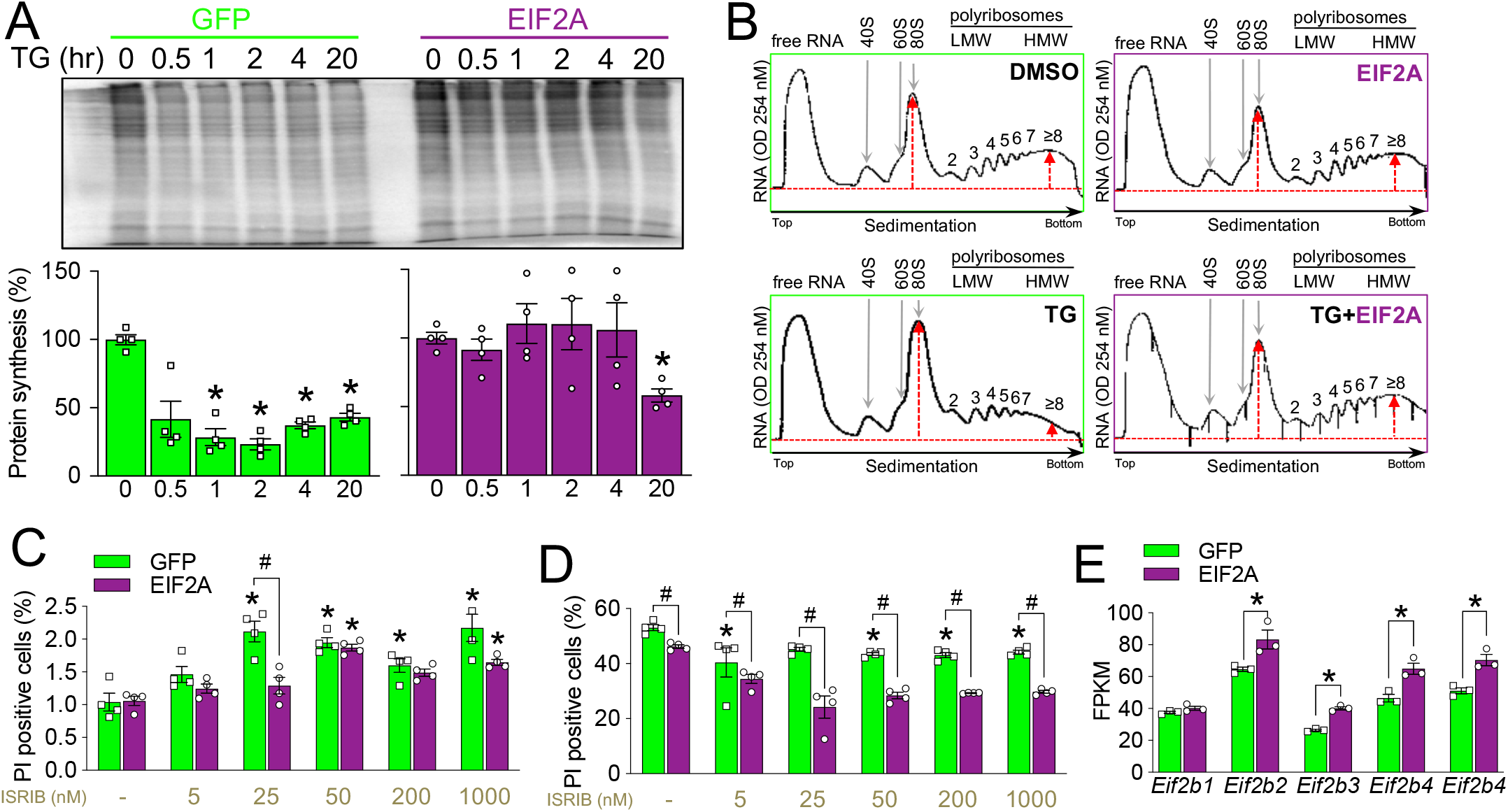
EIF2A over-expression prevents global translation attenuation during ER-stress. **(A)** Representative phosphor-images and densitometry quantification of total protein lysates from MIN6 cells overexpressing EIF2A:GFP or GFP and pulse-labeled with ^35^S-Met/Cys in the presence or absence of 1 μM thapsigargin (TG) for 0, 0.5, 1, 2, 4 or 20 hours. **(B)** RNA content measured at 254 nm in lysates from EIF2A:GFP or GFP overexpressing MIN6 cells, that were treated with 0.1% DMSO or 1 μM thapsigargin (TG) for 1 hour and fractionated on sucrose gradient into free RNA, 40S or 60S subunits, 80S monoribsomes, and low molecular weight (LMW) or high molecular weight polyribosomes (HMW). Numbers above peaks indicate number of ribosomes per mRNA. **(C**,**D)** Quantification of cell death assay using propidium iodide (PI) in MIN6 cells overexpressing EIF2A or GFP and treated with 0.1% DMSO **(C)** or with thapsigargin **(D)** for 24 hours in the absence or presence of 5 nM, 25 nM, 50 nM, 200 nM or 1000 nM ISRIB. **(E)** mRNA expression of *Eif2b* subunits in MIN6 cells overexpressing EIF2A compared to control MIN6 cells expressing GFP. *p<0.05

There is one known mechanism for bypassing the block on translation induced by phosphorylated EIF2S1: The small molecule ISRIB can override the phospho-EIF2S1-induced suppression at EIF2B. Specifically, ISRIB increases EIF2B protein abundance by preventing its proteasomal degradation (Rabouw et al., 2019). Thus, we compared the effects of multiple doses of ISRIB to EIF2A over-expression on β-cell death. In basal conditions, ISRIB was associated with a significant increase in β-cell death (Fig. 6C). At the 50 nM dose of ISRIB, this effect could be neutralized by EIF2A overexpression. In ER stress conditions induced by 1 μM thapsigargin, ISRIB treatment led to significant protection (Fig. 6D). Importantly, EIF2A overexpression further protect β-cells under these conditions. These additive effects, even at maximal ISRIB concentrations, strongly suggest that EIF2A has independent molecular mechanisms. Recent studies showed that ISRIB acts as a molecular scaffold, holding together eIF2B subunits along the assembly path to a fully active enzyme (Anand and Walter, 2020). Our RNAseq data revealed that mRNA levels of 4 out 5 *Eif2b* subunits were elevated in EIF2A over-expressing MIN6 cells (Fig. 6E), suggesting possible explanation for EIF2A-driven prevention of thapsigargin-induced translational inhibition.

## DISCUSSION

The goal of the present study was to determine the role of the novel translation initiation factor EIF2A in β cell surival. We found that EIF2A is relatively specific to islets, is upregulated by ER stress, and can be compensated for by EIF2D. We found that EIF2A overexpression protects rodent and human β cells from ER stress-induced apoptosis, *in vitro* and *in vivo*. EIF2A over-expression could overide phospho-EIF2S1-induced down-regulation of global protein synthesis, providing novel evidence that UPR control of translation can be uncoupled from apoptosis.

EIF2A exhibits high evolutionary conservation in eukaryotes, yet its role in the integrated stress response remains enigmatic. In *S. cerevisae* yeast, EIF2A is redundant for cell proliferation or survival, but mediates translation at internal ribosome entry sites (Reineke et al., 2011; Zoll et al., 2002). In mammalian cells, EIF2A helps viral mRNAs to avoid translation inhibition, when double-stranded RNA-activated EIF2AK2 phosphorylates EIF2S1 to block replication of Sindbis (Ventoso et al., 2006) or hepatitis C viruses (Kim et al., 2011). In recent years, several studies have examined the physiological role of EIF2A in various tissues (Golovko et al., 2016; Liang et al., 2014; Sendoel et al., 2017; Starck et al., 2012; Starck et al., 2016). Golovko and colleagues showed that whole pancreas lysates had the highest amount of EIF2A protein relative to other tissues tested (Golovko et al., 2016). We confirmed and extended these observations, demonstrating that EIF2A is highly abundant in pancreatic islets. Under basal conditions, EIF2A was localized primarily in the endoplasmic reticulum of β cells, in contrast to a human liver cell line where EIF2A is localized in the nucleus under normal conditions and is translocated to the cytoplasm under stress (Kim et al., 2011).

In our studies of mouse and human β-cells, EIF2A expression was increased in response to ER stress induced by thapsigargin or free fatty acid palmitate, as well as in low glucose conditions. Up-regulation of EIF2A has also been observed during integrated stress response in HeLa cells treated with bacterial subtilase cytotoxin and in primary mouse dendritic cells treated with various activators of EIF2AK2 (Starck et al., 2016). Our data, together with those of Starck and colleagues, clearly implicates EIF2A in the control of integrated stress response when EIF2S1 is phosphorylated and general protein synthesis is downregulated.

EIF2A appears to be part of a system with significant built-in redundancy. While a loss-of-function experiment in HeLa cells showed that EIF2A plays a non-redundant role in HSPA5 translation at steady-state and during induction of the ER stress with thapsigargin (Starck et al., 2016), we were unable to observe significant effects of partial EIF2A knockdown of HSPA5 protein levels in β cells. In the context of partial EIF2A knockdown using shRNA, we documented a concurrent compensatory increase in the expression of EIF2D, another alternative translation initiation factor able to initiate translation of some mRNAs regardless of EIF2S1 phosphorylation status (Skabkin et al., 2010). We found that EIF2D was not specific to β cells and, interestingly, EIF2D was significantly upregulated by palmitate but not thapsigargin. Cnop and colleagues reported that *EIF2D* mRNA was downregulated in palmitate-treated human islets by RNA sequencing (Cnop et al., 2013). Further biochemical studies to dissect the interplay between EIF2A, EIF2D and other possible alternative translation factors will be required and should ideally employ complete loss-of-function approaches. While global *Eif2a* knockout mice are generally viable (Golovko et al., 2016), the Mouse Genome Informatics resource lists “decreased circulating insulin level” as a phenotype in male *Eif2a*^em1(IMPC)Mbp^ strain of Eif2a mutant mice (MGI, 2020), consistent with our observation that EIF2A over-expression supports β cell survival and increased insulin secretion in the Akita mouse model of insulin misfolding stress.

Regulation of mRNA translation plays important role in the physiology and pathology of pancreatic β cells. On the one hand, β cells have an ability to ramp up insulin protein synthesis in response to elevated glucose levels via mechanisms that involve EIF2B (Gilligan et al., 1996). On the other hand, β cells have an ability to shut down mRNA translation to prevent their exhaustion. We previously provided the formal demonstration that insulin production itself is an inherent source of cellular stress in β cells (Szabat et al., 2016). Partial reduction of insulin production alleviated baseline ER stress, significantly modified ribosomal gene expression, and resulted in a paradoxical increase in the synthesis of non-insulin proteins (Szabat et al., 2016). Insulin makes up at least half of mRNA translation in β cells. Perhaps the focus on a single, hard-to-fold, secreted protein makes this cell type somewhat unique. Indeed, several drugs that have been shown to protect other cell types from ER stress, conversely have deleterious effects on β cells (Abdulkarim et al., 2017), suggesting fundamental differences in molecular mechanisms.

In the present study, we made the surprising observation that EIF2A over-expression circumvented the UPR-induced inhibition of global protein translation, despite sustained phosphorylation of EIF2S1. EIF2S1 evolved as an adaptive response to prevent accumulation of unfolded proteins in the ER and to reprogram mRNA translation rapidly to activate a cellular stress response. Attenuation of translation initiation by phosphorylation of EIF2SI is indispensable in survival of β cells but not of other cell types (Back et al., 2009; Harding et al., 2001; Scheuner et al., 2005). EIF2S1 phosphorylation functions as a switch during ER stress, causing inhibition of general mRNA translation initiation while promoting the translation initiation of specific stress-responsive mRNAs (Evans-Molina et al., 2013). It remains to be investigated whether the effects of EIF2A overexpression leads to preferential translation of a subset of mRNAs important for cell survival during ER stress, perhaps while the block on proteins that are specifically stress-inducing, such as insulin, remains. Interestingly, baseline ER stress in MIN6 β cells was increased by ISRIB, a drug that promotes EIF2B accumulation and can over-ride phospho-EIF2S1 inhibition of translation (Rabouw et al., 2019). Given that ISRIB is anti-apoptotic in many cell types, this surprising observation further supports the concept that β cells have relatively unique mechanisms controlling ER stress and translation. Under ER stress conditions, EIF2A overexpression circumvented the block on translation in the presence of phosphorylated EIF2S1, something only ISRIB has previously been shown to do. We reasoned that combining ISRIB with EIF2A overexpression would allow us to determine whether these manipulations have independent mechanisms of action. Indeed, the protective effects of EIF2A overexpression were additive to those of ISRIB at all doses tested. Our interpretation of these data is that EIF2A overexpression protects β cells via a novel mechanism independent of the blockage of EIF2B degradation induced by ISRIB. Indeed, RNA sequencing revealed upregulation of multiple EIF2B subunits mRNAs with EIF2A overexpression. Perhaps EIF2A increases EIF2B protein by increasing transcription, while ISRIB prevent EIF2B degradation. It is also possible that EIF2A acts completely independently of the EIF2 complex to recruit Met-tRNA_i_ to the 40S ribosomal subunit. RNA sequencing identified additional potential mediators of β cell-protective effects of EIF2A, including up-regulated transcripts (*Ins1, Sct, Malat1, Basp1, Cdhr5, Ano1*) and down-regulated transcripts (*Spp1, Nr4a1, Cldn11, Aldh1a3, Pdzd2*). Cleary multiple mechanisms are involved β cell protection by EIF2A. Taken together, our study defines new roles for the alternative translation initiation factor EIF2A during β cell ER stress, integrating the UPR, protein synthesis, and cell fate decisions.

## METHODS

### Cell culture

MIN6 and HEK293T cells were cultured in Dulbecco’s modified eagle’s medium (Thermo Fisher Scientific) containing 22.2 mM glucose, 100 units/ml penicillin, 100 mg/ml streptomycin and 10% (v/v) FBS. Mouse pancreatic islet were isolated from 12-to 20-week-old male C57BL/6J mice (Jax, Bar Harbor, MA, USA) using collagenase and filtration. All experiments using animals were approved by the UBC Animal Care Committee in accordance with national guidelines and international standards. The islets were further handpicked using a brightfield microscope. Islets were cultured overnight (37°C, 5% CO2) in RPMI1640 medium (Thermo Fisher Scientific) supplemented with 100 units/ml penicillin, 100 mg/ml streptomycin (Thermo Fisher Scientific) and 10% (v/v) FBS (Thermo Fisher Scientific). Human islets, obtained under informed consent for the use of pancreatic tissue in research, were provided by the Ike Barber Human Islet Transplant Laboratory (University of British Columbia, Vancouver) and by the Alberta Diabetes Institute IsletCore (University of Alberta, Edmonton). Protocols approved by the University of British Columbia Research Ethics Board and cultured under the same conditions as mouse islets.

### Gene expression, transcriptomics, and network analysis

RNA was isolated from cell or islet samples using RNeasy mini kit (Qiagen, #74106) according to manufacturer’s instructions. cDNA was synthesized using qScript™ cDNA synthesis kit (QuantaBio, #95047-500). qPCR was performed in using StepOnePlus Real-Time PCR System (Applied Biosystems).

RNA Sequencing was performed at the UBC Biomedical Research Centre Sequencing Core. Briefly, sample quality control was performed using the Agilent 2100 Bioanalyzer. Qualifying samples were then prepped following the standard protocol for the NEBnext Ultra ii Stranded mRNA (New England Biolabs). Sequencing was performed on the Illumina NextSeq 500 with Paired End 42bp × 42bp reads. De-multiplexed read sequences were then aligned to the reference sequence using STAR (https://www.ncbi.nlm.nih.gov/pubmed/23104886) aligners. Assembly were estimated using Cufflinks (http://cole-trapnell-lab.github.io/cufflinks/) through bioinformatics apps available on Illumina Sequence Hub.

Gene differential expression was analyzed using DESeq2 R package (Love et al., 2014). KEGG pathways were enriched from all differentially expressed genes using clusterProfiler R package (Yu et al., 2012). All the R scripts are available on GitHub repository https://github.com/hcen/EIF2A. Protein-protein interaction network and transcription factor-gene network were generated by online tool NetworkAnalyst3.0 (www.networkanalyst.ca) using default settings (Zhou et al., 2019). Specifically, the protein-protein interaction network was based on STRING interactome with high (900) confidence score and experimental evidence. ENCODE database was selected to generate the transcription factor-gene network. Transcription factors that were not expressed in our RNAseq were filtered out.

EIF2A and EIF2D interacting proteins were predicted with STRING using the high (900) confidence score and experimental evidence settings.

### Immunoblot and co-immunoprecipitation

MIN6 cells, mouse or human pancreatic islets were homogenized and sonicated in RIPA buffer (50 mM β-glycerol phosphate, 10 mM HEPES, 1% Triton X-100, 70 mM NaCl, 2 mM EGTA, 1 mM Na_3_VO_4_, and 1 mM NaF) supplemented with complete mini protease inhibitor cocktail (Roche, Laval, QC). Protein lysates were resolved by SDS-PAGE and transferred to PVDF membranes (BioRad, CA) and probed with antibodies against, p-EIF2S1 (Ser51) (1:1000, Cat. # 3398), EIF2S1 (1:1000, Cat. #5324), HSPA5 (1:1000, Cat. #3183), cleaved-caspase 3 (1:1000, Cat. #9664), ERN1 (1:1000, Cat. #3294), EIF1A (1:1000, Cat. # #12496), EIF5A (1:1000, Cat. #20765) all from Cell Signalling (CST), and EIF2A (1:1000, Cat. # 11233-1-AP, ProteinTech), EIF2D (1:1000, Cat. #12840-1-AP, ProteinTech), DDIT3 (1:1000, #MA1-250, ThermoFischer), β-tubulin (1:1000, Cat. #T0198, Sigma). The signals were detected by secondary HRP-conjugated antibodies (Anti-mouse, Cat. #7076; Anti-rabbit, Cat. #7074; CST) and Pierce ECL Western Blotting Substrate (Thermo Fisher Scientific). Protein band intensities were quantified with Adobe Photoshop sotware.

### Immunofluorescence staining

Pancreata were perfused, then fixed for 24 hours with 4% paraformaldehyde, and then washed twice with 70% ethanol prior to paraffin embedding and sectioning (5 μm) to obtain 5 different regions of the pancreas (100 μm apart) by WAXit (Vancouver, Canada). Paraffin was removed by 5 min xylene incubation steps. Sections were rehydrated in decreasing concentrations of ethanol and rinsed with water and PBS. Epitope retrieval was done either by immersing samples 10 mM citrate buffer, pH 6.0 for 15min at 95°C, or by transferring sections to prewarmed 1N HCl for 25 min at 37°C. Samples were washed with PBS twice and were either blocked for 10 min at room temperature (Dako protein block #X0909), or with goat block (GB) with Triton X-100 (10% BSA + 5% Goat Serum with 0.5% Triton X-100) for 1-4 hours at room temperature. Samples were incubated overnight at 4°C in primary antibodies targeting anti-insulin (1:200, Abcam #Ab7872, Dako #A0564), anti-glucagon (1:100, CST, #2760S), anti-HSPA5 (1:100, CST #3183), anti-DDIT3 (1:100, ThermoFischer #MA1-250) Following 3 PBS washes (5min each), samples were incubated for 30min or 1h at room temperature in secondary antibodies in a light-deprived humid chamber. Secondary antibodies applied were anti-rabbitAlexa Fluor-488 (1:200, Invitrogen, # A-11008), anti-rabbit Alexa-488 (1:200, Invitrogen, #A11034), anti-rat Alexa-594 (1:200, Invitrogen, #A11007), anti-guinea pig Alexa-647 (1:200, Invitrogen, #A21450), anti-guinea pig Alexa-594 (1:200, Invitrogen #A-11076). Samples were mounted with either VECTASHIELD Hard Set Mounting Medium (Vector labs, # H-1500) following an additional three washes in PBS (10min each). β cell area was calculated as insulin positive area normalized to the entire pancreas of each section using Zeiss 200M microscope.

### Live cell imaging

MIN6 cells or dispersed human islets were seeded in 96-well plates and cultured for 24hr. Cells were stained with Hoechst 33342 (Sigma-Aldrich) and propidium iodide (Sigma-Aldrich), prior to imaging with ImageXpress^MICRO^ (Molecular Devices, Sunnyvale, CA, USA) as previously described (Yang and Johnson, 2013).

### Analysis of total protein synthesis rate

For pulse labeling of newly translated proteins MIN6 cells were incubated in complete DMEM media but without cysteine and methionine (MP Biomedicals, #SKU 091646454) for 1hr. Subsequently media was supplemented with 250 μCi of [^35^S]-cysteine/methionine mixture (PerkinElmer, NEG772002MC) and cells incubated under normal conditions for 30 min. Cells were then lysed and proteins separated by SDS-gel electrophoresis as described above. Gels were fixed for 30 min in 50% (v/v) ethanol in water with 10% (v/v) acetic acid, dried in gel dryer (Bio-Rad model 583) and then exposed to storage phosphor screen (GE Healthcare) overnight. Screens were imaged and digitized using Typhoon FLA 9000 biomolecular imager (GE Healthcare). Protein bands intensities were quantified with Adobe Photoshop software.

### Polysome fractionation

Ribosome fractionation using sucrose density gradient centrifugation was performed as previously described (Gandin et al., 2014).

### AAV6 vector construction, viral production and in vivo administration

The single-stranded AAV vector plasmid (pssAAV) expressing EIF2A under control of rat *Ins2* gene promoter (RIP) was made using existing pssAAV-RIP by inserting SalI-EIF2A-GFP-NdeI fragment, which was obtained by PCR from plasmid pCMV6-EIF2A-GFP (MG209105, OriGene, Rockville, MD) using forward primer 5’-ATTCGTCGACTGGATCCGGT-3’, and a reverse primer 5’-CACACATATGTTAAACTCTTTCTTCACCGGCA-3’. pssAAV expressing GFP under control of rat *Ins2* was made by deleting the EIF2A open reading frame. AAV6 viral particles were produced and purified by SignaGen Laboratories (Rockville, MD).

C57BL/6-*Ins2*^Akita^/J (003548) 5-week-old female mice were purchased from The Jackson Laboratory (Bar Harbor, Maine). Viral delivery to the pancreas was performed via intraductal injection of the 6-week-old mice randomized into two groups (2 cohorts, n=10-11), as previously described (Obach et al., 2018). Briefly, after exposing the common bile duct, one microclamp was placed on it to close to the gall bladder. Another microclamp was placed under the sphincter of Oddi to avoid leakage of the vector into the duodenum. Then 1.5 × 10^11 viral particles in 100 μl of PBS were slowly injected into the pancreatic duct via sphincter of Oddi through 30 G needle attached to the 1 ml syringe. The needle entry point was sealed with 3M Vetbond adhesive and clamps were removed.

### Blood collection and in vivo analysis of glucose homeostasis and insulin secretion

Tail blood was collected for blood glucose measurements using a glucometer (One Touch Ultra 2 Glucometer, Lifescan, Canada) for single time points as well as during glucose and insulin tolerance tests. Mice were fasted for 6h during glucose and insulin tolerance tests and glucose stimulated insulin secretion tests. For intraperitoneal (*i*.*p*.) glucose tolerance tests, the glucose dose was 1 g/kg. Femoral blood was collected for single timepoints, as well as for measurements of *in vivo* glucose-stimulated insulin secretion after *i*.*p*. injection of 1 g/kg glucose. Blood samples were kept on ice during collection, centrifuged at 2000rpm for 10min at 4°C and stored as plasma at -20°C. Plasma samples were analysed for insulin (Stellux Chemi Rodent Insulin ELISA, Alpco #80-INSMR-CH10) and proinsulin (Alpco #80-PINMS-E01). Measurements were performed on a Spark plate reader (TECAN), and analysed using GraphPadPrism (GraphPad Software Inc., San Diego, CA).

### Islet isolation and dispersion

Mouse islet isolations were conducted by ductal inflation and incubation with collagenase, followed by filtration and hand-picking as in our previous studies and following a protocol adapted from Salvalaggio (Luciani and Johnson, 2005). 24h post islets isolations, islets were washed (x4) (Ca^2+/^Mg^2+^-Free Minimal Essential Medium (Gibco) followed by gentle trypsinization (0.01%), and resuspended in RPMI 1640 (Gibco) supplemented with 10%FBS, 1%P/S. Seedings were done according to the experimental procedure.

### Statistics

Data were analyzed in GraphPad Prism using a one-way multiple comparisons ANOVA followed by a *post hoc* t-test using the Tukey HSD method. Data are expressed as means ± SEM, and p-value less than 0.05 was considered significant.

## ACKNOWLEDGEMENTS

We thank members of the Johnson and Jan labs for helpful discussions.

